# Bessel Beam Optical Coherence Microscopy Enables Multiscale Assessment of Cerebrovascular Network Morphology and Function

**DOI:** 10.1101/2024.04.16.589730

**Authors:** Lukas Glandorf, Bastian Wittmann, Jeanne Droux, Chaim Glück, Bruno Weber, Susanne Wegener, Mohamad El Amki, Rainer Leitgeb, Bjoern Menze, Daniel Razansky

## Abstract

Understanding the morphology and function of large-scale cerebrovascular networks is crucial for studying brain health and disease. However, reconciling the demands for imaging on a broad scale with the precision of high-resolution volumetric microscopy has been a persistent challenge. In this study, we introduce Bessel beam optical coherence microscopy with an extended focus to capture the full cortical vascular hierarchy in mice over 1000 × 1000 × 360 μm^3^ field-of-view at capillary level resolution. The post-processing pipeline leverages a supervised deep learning approach for precise 3D segmentation of high-resolution angiograms, hence permitting reliable examination of microvascular structures at multiple spatial scales. Coupled with high-sensitivity Doppler optical coherence tomography, our method enables the computation of both axial and transverse blood velocity components as well as vessel-specific blood flow direction, facilitating a detailed assessment of morpho-functional characteristics across all vessel dimensions. Through graph-based analysis, we deliver insights into vascular connectivity, all the way from individual capillaries to broader network interactions, a task traditionally challenging for *in vivo* studies. The new imaging and analysis framework extends the frontiers of research into cerebrovascular function and neurovascular pathologies.

## INTRODUCTION

Cerebral vasculature, extending from large pial vessels to the capillary bed, constitutes a complex network fundamental to supplying blood to the cortex^1^. Blood circulation facilitates crucial brain functions via oxygen and glucose delivery, whilst its failure plays a crucial role in brain pathologies such as Alzheimer’s or stroke^2,3^. Ample blood supply to all tissue regions is governed by the complex interplay of the vascular network structure and blood flow^4,5^. It is thus imperative to comprehensively investigate vessel morphology and function across multiple vascular scales, from the microscopic capillary dimensions to the network relationships on a macroscopic scale. Due to extensive interconnectivity of the vasculature, many pathologies result in long- and short-term blood flow velocity alterations. For example, Alzheimer’s has been linked with long-term cortical hypoperfusion, affecting vessel diameters as well as blood flow velocities^6,7^. In stroke, a sudden local hypoperfusion is caused by obstruction of a major vessel, but blood flow may even increase in the surrounding regions in an attempt to compensate through the remaining vascular network^8^. Quantifying these functional regulations is critical for a better understanding and treating neurovascular pathologies, which can be greatly facilitated by joint evaluation of morphological and functional flow repercussions *in vivo*.

Intravital imaging techniques, such as laser speckle contrast imaging (LSCI) and two-photon microscopy (2PM), are widely employed in pre-clinical neurovascular research. However, these techniques fall short of capturing the complexity of vascular networks and their variable flow dynamics across multiple scales. Recently, ultra high-speed 2PM approaches have been developed^9^, yet no large-scale velocimetry assessments were reported. Extended-focus 2PM systems, often employing Bessel beams, increase the number of resolved vessels to the detriment of depth resolution^10^. The dynamic light scattering imaging technique^11^ can improve upon the spatial resolution limitations of LSCI while addressing its other drawbacks, such as overestimation of blood perfusion reduction due to stroke. But the approach remains inherently two-dimensional and lacks capillary resolution. Other imaging techniques have also been successfully used for functional brain studies in health and disease. Optoacoustic tomography can resolve vascular morphology and oxygenation in the whole mouse brain at high frame rates but its spatial resolution is only suitable for rendering large brain vessels. Its high-resolution optoacoustic microscopy counterpart sacrifices imaging speed and/or field-of-view (FOV) to attain capillary level resolution in the lateral plane, yet it provides poor axial resolution that hinders accurate depiction of three-dimensional microvascular networks^12,13^. Recently, localization-based approaches, such as ultrasound localization microscopy (ULM)^14^, localization optoacoustic tomography (LOT)^15^ and widefield fluorescence localization microscopy (WFLM)^16^ are gaining interest due to their high spatial resolution as well as intrinsic velocity measurements across large FOVs. However, ULM and LOT are not yet able to resolve the capillary bed whilst WFLM is only suitable for superficial investigations further lacking depth resolution. All in all, none of the currently available approaches can simultaneously provide vessel morphology and blood velocity measurements at isotropic capillary-level 3D resolution across a large (millimeter-scale) FOV.

Optical coherence tomography (OCT) is a powerful technique for rapid, label-free, three-dimensional imaging of vascular morphology and function. When studying cortical vasculature in mice, optical coherence tomography angiography (OCTA) provides ample resolution to resolve even the smallest capillaries while covering larger volumes at faster scanning times in comparison to 2PM^17^. However, OCTA is plagued by image artifacts that often prohibit accurate volumetric blood vessel segmentation^18^. Nevertheless, recent deep learning segmentation approaches can adequately deal with OCTA artifacts and extract vascular networks, allowing vessel-wise quantitative analysis^7^. Yet, these approaches are currently unable to fully capture the cortical vasculature’s network characteristics, encompassing the blood flow from pial arteries into diving arterioles through the capillary bed and back through the venous tree. This is largely due to the hard compromises for the achievable FOV and depth-of-field (DOF) at the given capillary resolution levels. While depth scanning can be feasible, it prolongs imaging times thus diminishing OCTA’s advantages in comparison to 2PM.

Additionally, Doppler optical coherence tomography (DOCT) is capable of probing *in vivo* hemodynamics across the entire imaged volume. However, DOCT predominantly provides axial flow velocity information whilst its performance is usually compromised in slow-flowing capillaries^19,20^. Recent efforts have successfully focused on extending the Doppler signal detection range towards capillary level flows, as well as estimating transverse flow velocities in larger vessels^21,22^. Transverse blood flow velocity has also been estimated from OCTA decorrelation times that however may saturate in larger vessels and require carefully acquired calibration curves^23^. Alternatively, a direct detection of RBC passages in capillaries from the OCT signal constitutes an elegant way to deal with the discrete nature of blood flow thus rendering RBC flux akin to 2PM. But this approach is likewise limited to slow flows and smaller vessels^17^.

To overcome these limitations, we propose an integrated acquisition and analysis pipeline that yields comprehensive metrics for both blood flow velocities as well as morphological characteristics of the vascular network, all across vessel scales and with minimal manual intervention. Our approach capitalizes on the extended-focus optical coherence microscopy (OCM) concept, allowing to accurately capture vascular network connectivity without depth-scanning. The method enables a simultaneous, multiscale precision assessment of morphology and function, a crucial need in understanding and developing treatments for complex neurovascular conditions such as stroke or Alzheimer’s. Taking advantage of deep neural networks for segmenting OCTA volumes, the automatic pipeline facilitates quantitative analysis of large datasets that would otherwise be unfeasible. DOCT with unprecedented capillary flow sensitivity further provides axial and transverse blood flow velocities as well as flow direction. Finally, we present a microvascular network analysis, quantifying cerebral blood flow along its transport route from pial arteries, through the capillary bed and out via the pial vein structure.

## RESULTS

### Extended depth-of-field enables single snapshot visualization of vascular network parameters

Increasing the available FOV without sacrificing resolution is a highly sought-after feature in any microscope design. A typical approach to overcome the corresponding trade-offs is depth scanning. This helps maintaining high lateral resolution while preserving high resolving capacity in the axial dimension, but comes at the cost of substantially extended measurement time. In contrast, OCM inherently provides axial sectioning along the simultaneously recorded axial (depth) dimension, leading to hard trade-offs between the lateral resolution and confocally gated imaging depth. As a result, standard OCM systems typically lack the distinct advantage of high imaging speed as several depth-resolved yet confocally gated axial scans need to be stitched at multiple axial positions, which often renders other techniques like 2PM better suited for high resolution imaging of microvascular networks.

We alleviate this challenge by employing an extended-focus approach. Our extended-focus infrared OCM (xf-irOCM) system (Fig. 1a) is devised as a Mach-Zehnder interferometer and uses an axicon in the sample arm for Bessel beam synthesis^24^. In contrast to common Gaussian beams, Bessel beams provide a significantly expanded DOF for microscopy designs^10,25^. This becomes particularly attractive when paired with OCM, which does not rely on optical sectioning to render high axial resolution. In contrast to the previously reported designs, a more powerful light source and optimized beam-splitter ratios result in an over three-times increased light power at the sample while still providing ample reference arm power. This provides sufficient signal-to-noise ratio for Doppler imaging along the entire DOF.

**Figure 1:**
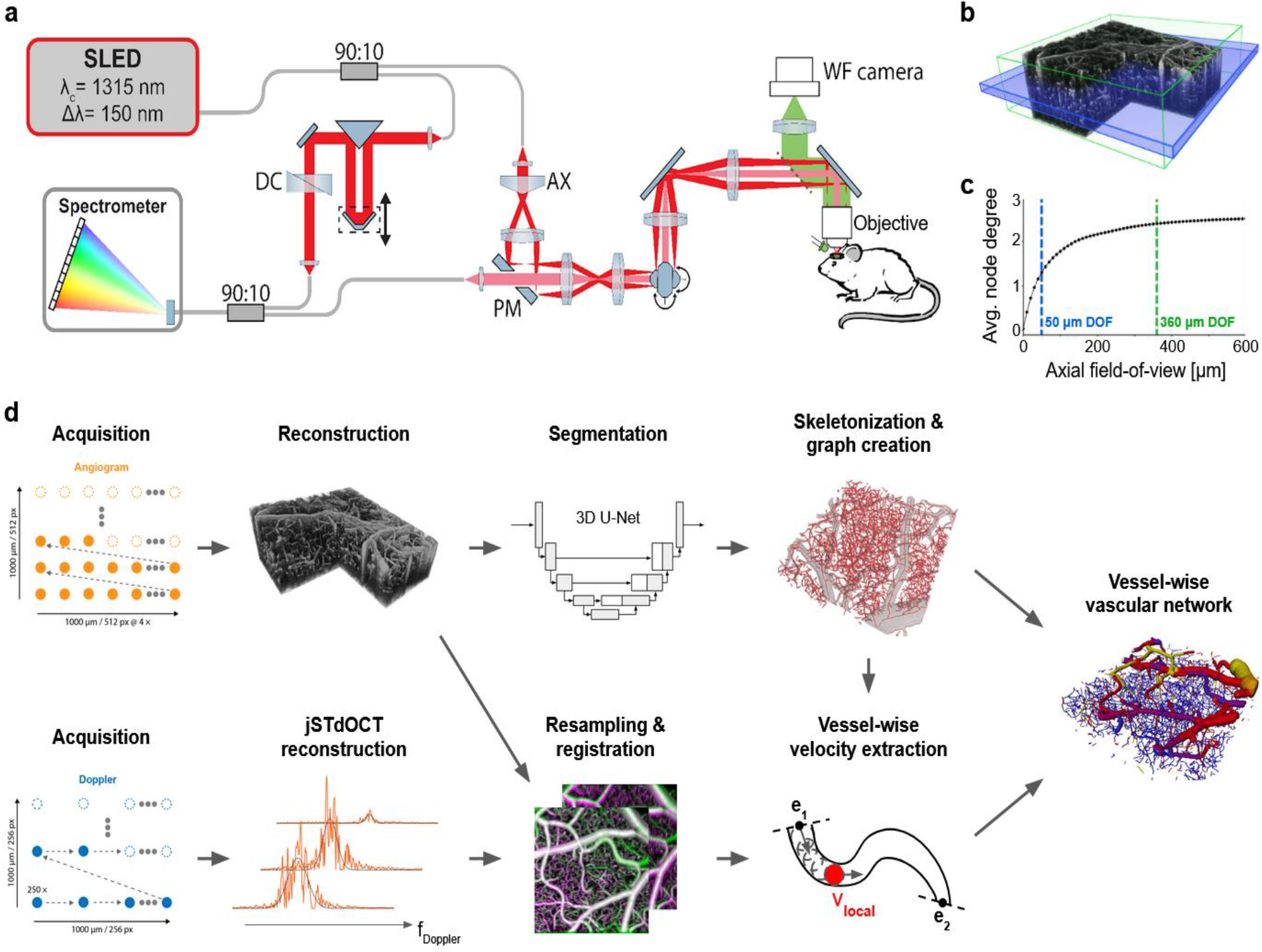
Overview of the xf-irOCM and processing pipeline. a) Schematic of the xf-irOCM. In the sample arm, illumination and detection light paths are decoupled using a pierced mirror (PM) and depicted in dark and light red, respectively. AX: axicon, DC: dispersion compensation b) OCTA volume acquired with the xf-irOCM. The green box emphasizes the in-focus volume achieved with our xf-irOCM, resulting in a depth-of-field of ≈360 μm. The blue box represents the simulated in-focus volume (≈50 μm) of a standard OCM system with equivalent lateral resolution. The lateral dimensions are 1000 μm × 1000 μm. c) Average node degree of the vascular networks from Blinder et al.^1^ over increasing axial field-of-view. These vascular networks serve as a reference to estimate the necessary axial FOV to duly capture vascular network characteristics. The dashed green and blue lines highlight the extended depth-of-field and regular depth-of-field shown in (b), respectively. d) Acquisition and automatic processing pipeline that enables extraction of vessel morphology and functional flow information in each vessel, combining OCTA and DOCT.

Fig. 1b showcases a volumetric angiogram recorded with our xf-irOCM. The green box represents the in-focus volume measuring 1000 × 1000 × 360 μm^3^. Capillaries are clearly resolved throughout the volume. In contrast, the blue box describes the DOF of a simulated OCM system that employs a traditional Gaussian beam and is otherwise identical to our system, achieving the same lateral resolution of 3 μm in tissue. Its DOF is limited to roughly 50 μm in the ideal simulation case. Hence, when compared to a standard confocal implementation, our xf-irOCM achieves a minimum 7-fold increase in the acquired volume of image data within the same acquisition time. To further quantify the advantage of the xf-irOCM when analyzing microvascular networks, we highlight its ability to accurately capture large-scale network characteristics. In this context, we use a previously published dataset that provides high-quality vascular network segmentation maps in graph format acquired using *ex vivo* imaging techiques^1,26^. From these segmentation maps we extract the average node degree (Fig. 1c), where each node represents a blood vessel bifurcation or high curvature point, over increasing depth range. The resulting curve emphasizes that narrow axial FOVs sever many vascular network interconnections with the average node degree approaching an almost constant value of approximately 2.57 for increased axial FOV. In comparison, the Gaussian system DOF (blue line) majorly underestimates the average node degree to be 1.31, which is equivalent to 49.2 % of the steady-state value of 2.57, because many capillary connections are severed by the limited axial FOV. In contrast, the axial extent that is equivalent to the xf-irOCM’s DOF (green line) captures vessel connections with a much smaller 6.7 % error percentage by estimating the average node degree to be 2.39. Therefore, no axial scanning is necessary to obtain accurate vascular network connectivity when imaging with the xf-irOCM.

### Volumetric total flow velocity measurements across vessel scales using highly sensitive Doppler OCT and extended-focus OCM

An overview of the integrated processing pipeline is presented in Fig. 1d. Angiograms and Doppler datasets are acquired sequentially but without idle time. Voxel-wise Doppler spectra are obtained using the joint spectral and time domain OCT (jSTdOCT) framework^27^. Additionally, two crucial steps have been added. First, high spatio-temporal oversampling of 250 A-scans at each position enables detection of slow-flowing capillaries by yielding a highly resolved Doppler spectrum and narrow constant signal peak at zero frequency, referred to as DC peak in the following. Second, high-pass filtering along the time domain allows suppression of the DC peak in the Doppler spectra, arising from mixed signals within voxels. Without the filter, the signal from slow-flowing capillaries is strongly overshadowed by the DC peak. The Doppler power spectra (DPS) are then obtained by Fourier-transforming all A-scans at each lateral position along the time domain. Then, a modified Gaussian function is fitted to the resulting DPS. Subsequently, axial and transverse velocity components are calculated from the DPS’s mean and standard deviation, as previously derived for Bessel beam OCM systems^28^. Taken from an exemplary high-resolution angiogram (Fig. 2a), the depth slice in Fig. 2b serves as an example for flow velocity measurements across vessel scales. The Doppler shift and Doppler standard deviation maps from the same slice are presented in Fig. 2c & 2d, respectively. The red arrow in the angiogram labels the cross-section of an artery. Its DPS (Fig. 2e) shows a Doppler shift μ of 2119 Hz that is obtained by fitting a modified Gaussian to the orange signal, solely depending on the axial velocity component. This is equivalent to the Doppler shift that is commonly calculated in DOCT^20^. However, in contrast to the standard autocorrelation approach, the fit allows calculating the DPS’s standard deviationσ that widens with increased transverse flow velocities thus allowing their estimation^22,28,29^. For the artery,σ is calculated to be 3817 Hz, resulting in transverse flow velocity of 12.2 mm/s, which is higher than the vein’s transverse flow velocity of 8.5 mm/s (blue arrow in Fig. 2b, spectrum Fig. 2 f).

**Figure 2:**
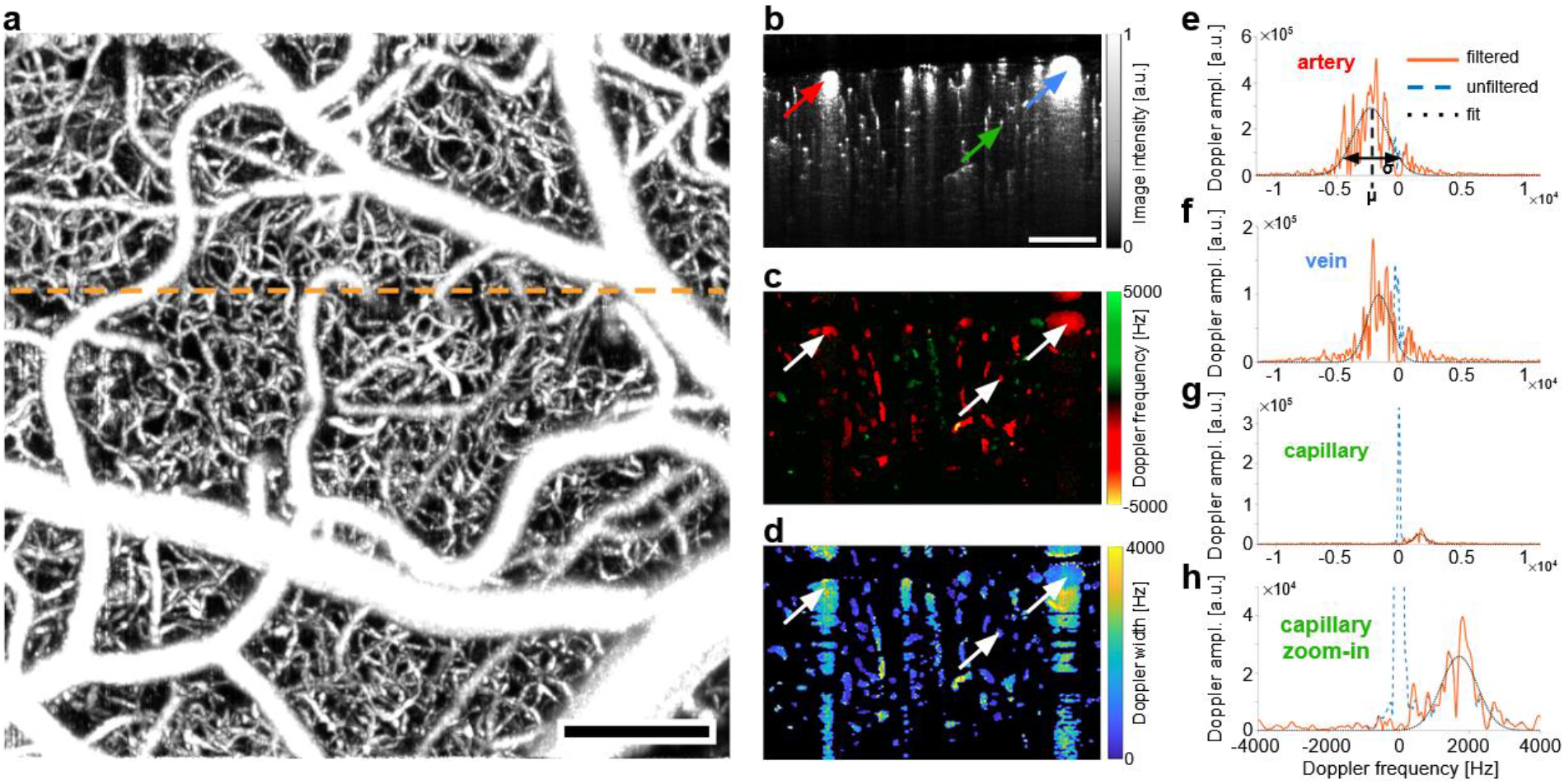
Rendering volumetric flow velocity maps using highly sensitive DOCT. a) Exemplary maximum intensity projection of an OCT angiogram. The dashed orange line indicates the slice position of b-d. Scalebar: 200 μm. b) Angiogram depth slice. The red, blue and green arrows mark the exemplary artery, vein and capillary in e-h, respectively. c) Doppler shift depth slice. The white arrows highlight the vessels in b) at the same positions. d) Doppler standard deviation depth slice. e-h) Doppler power spectra from the voxels indicated in b). The high-pass filtered DPS is shown in orange, the raw DPS as the dashed blue curve and the resulting modified Gaussian fit as the dotted black line. The Doppler frequency shift μ is calculated as the fit’s peak position and the Doppler frequency spread σ as its standard deviation.

For the case of comparatively wide arteries and veins, differences between the high-pass filtered signal (orange) and unfiltered signal (blue) are minimal and do not significantly affect the fit. A different perspective emerges when examining the DPS (Fig. 2g) from thin and slow-flowing capillaries (green arrow in Fig. 2b) that make up the bulk of the cerebral vasculature. As capillary diameters approach the voxel size, a blending of static and dynamic components occurs^30,31^. In this scenario, the DC peak of the unfiltered DPS dwarfs the capillary signal of interest, as illustrated in a zoomed-in view (Fig. 2h). Nevertheless, after high-pass filtering, a good fit can be achieved with the axial and transverse flow velocities accurately extracted from the capillary. The additional flow phantom experiments reaffirmed accuracy of the flow velocity estimations (Fig. S1 & S2).

### Deep learning-based vessel segmentation facilitates network wide mapping of blood flow velocity and direction

Analyzing vascular networks across scales can be accomplished by transforming the information from the image domain into a graph-based representation^1^. However, such transformation necessitates accurate segmentation of the imaged vasculature. Although OCTA can accurately resolve capillaries, the data is often afflicted with tail artifacts, locally varying SNR, and signal intensity loss with vessel orientation, making the segmentation and accurate delineation of the axial blood vessel extent difficult when using traditional methods such as thresholding. Numerous strategies aimed at alleviating tail artifacts, such as shape filtering^32^ and g1-autocorrelation^33^, have not yielded satisfactory performance and fast acquisitions without suppressing some capillary structures. While tail artifacts are less pronounced in Bessel beam OCM due to self-healing beam properties^34,35^, the existing thresholding approaches remain inadequate^36^.

Instead, we utilize a supervised deep learning-based approach that allows for accurate segmentation of vasculature across all scales in the murine cortex, from large pial vessels to capillaries. Specifically, we finetune a pre-trained 3D U-Net^37–39^ on a 350 × 350 × 350 μm^3^ manually annotated volume. Pre-training was performed on simulated 3D data, relying on synthetic vessel trees constructed from physiological principles^40^. We observe that pre-training, to some extent, reduces the need for extensive manual annotation of 3D OCT angiograms. Exemplary segmentation maps inferred by the trained 3D U-Net are depicted in Fig. 3a & 3b. Analysis of the predicted segmentation maps leads to the conclusion that we manage to successfully eliminate tail artifacts while capillaries are accurately segmented even at the smallest scale and throughout the entire FOV.

**Figure 3:**
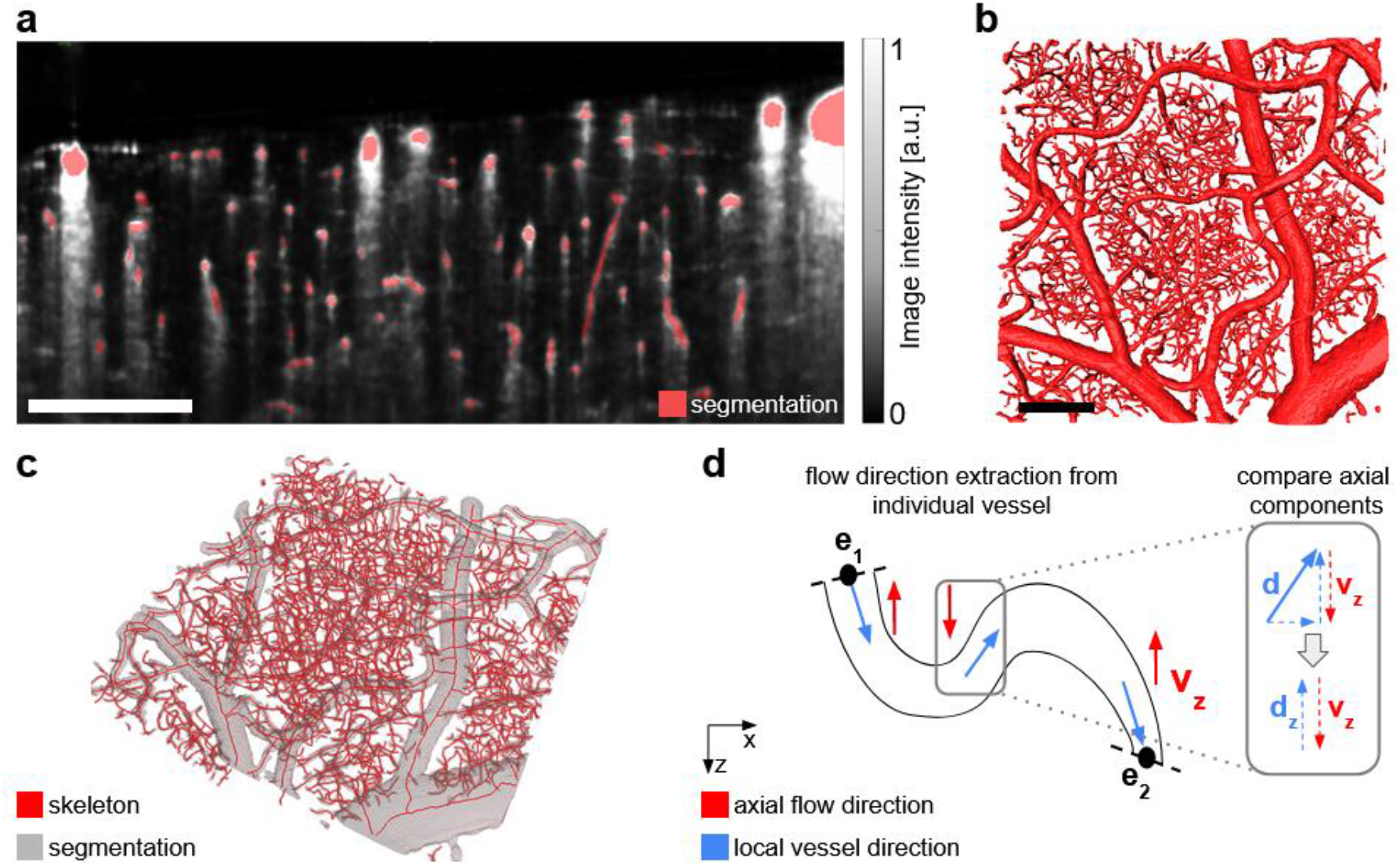
Segmentation and flow velocities. a) Side-view of the volumetric vessel segmentation map (red), overlayed onto the underlying OCTA slice (gray). b) Top-view of the volumetric vessel segmentation output from the neural network. c) Extracted vessel skeleton and the underlying vessel segmentation map after cleaning. d) Principle of flow direction extraction. An exemplary vessel is shown as its y-axis projection. Blue arrows describe the arbitrarily chosen direction along the vessel skeleton from one endpoint (e_1_) to the second endpoint (e_2_). Red arrows describe the local axial flow in z-projection that is obtained from the standard Doppler signal. At each skeleton voxel, the axial components of the 3D vessel skeleton and the flow velocity are compared (inlet). All scalebars are 200 μm.

The predicted segmentation map is subsequently preprocessed by employing morphological closing for smoothing (grey, Fig. 3c). By leveraging a skeletonization algorithm optimized for vasculature^41^, smooth vessel centerlines are obtained (red, Fig. 3c). From this, a vascular network graph is derived in which vessel bifurcation points and their connecting vessel segments are respectively represented as nodes and edges. Next, flow velocities are extracted for each vessel segment. Because Doppler and angiogram acquisitions are performed sequentially at the same position, axial and total blood flow velocities can be extracted along each vessel segment through local averaging after registering the datasets to correct for the differing scanning protocols and upscaling them to the same spatial resolution level.

Blood flow velocities are not only characterized by their magnitude but also by their direction. DOCT is able to measure the axial component of the flow, including its direction, relative to the probe beam. We show that combining the spatial constraints of the 3D vessel segmentation maps with the extracted axial flow component allows to retrieve the blood flow direction in all vessels that exhibit detectable axial flow (Fig. 3d). By arbitrarily defining a direction along the vessel centerline (blue arrows) and comparing the local axial orientation of the vessel segment skeleton with the Doppler flow direction (red) at each position, the respective blood flow direction for each vessel can readily be determined. Because tortuous capillaries can be partially or fully oriented perpendicular to the imaging direction, few vessels may have no Doppler signal. In such cases, we take advantage of the determined flow velocities and directions in the directly interconnected surrounding vessels. By enforcing zero net flow at each bifurcation point, the missing flow velocity and direction can be recovered if a vessel is connected to a bifurcation point with otherwise known blood flow velocities and flow directions (Fig. S5). Typically, less than 8 % of vessel segments cannot be assigned a velocity. After correction, this number drops below 2 %.

### Known flow directions enable artery/vein classification of pial vasculature and determination of the network-wide branching order

The resulting microvascular network and total blood flow velocities are depicted in Fig. 4a. The vessel diameter distribution (Fig. 4b) agrees well with previously reported data^26,42^. Furthermore, the imaged microvasculature exhibits a velocity distribution that peaks between 1 – 3 mm/s with >85 % of vessel having flow velocities below 5 mm/s (Fig. 4c). These are almost exclusively found in the capillary bed having vessel diameters below 8 μm. Higher velocities are predominantly manifested in larger vessels, as also evident from the volumetric renderings (Fig. 4a). These vessel segments exhibit longer lengths between branching points, resulting in lower histogram counts for high velocities. To account for these effects, we furthermore examined flow velocities independently for each vessel type, namely, the arteries, veins and capillaries, and relative to their perfused volume (Fig. 4d). Pial arteries exhibit the highest flow velocities, reaching up to 30 mm/s, while accounting for 21 % of the total blood volume. In comparison, pial veins show reduced blood flow velocities but a larger perfused blood volume of 33 %.

**Figure 4:**
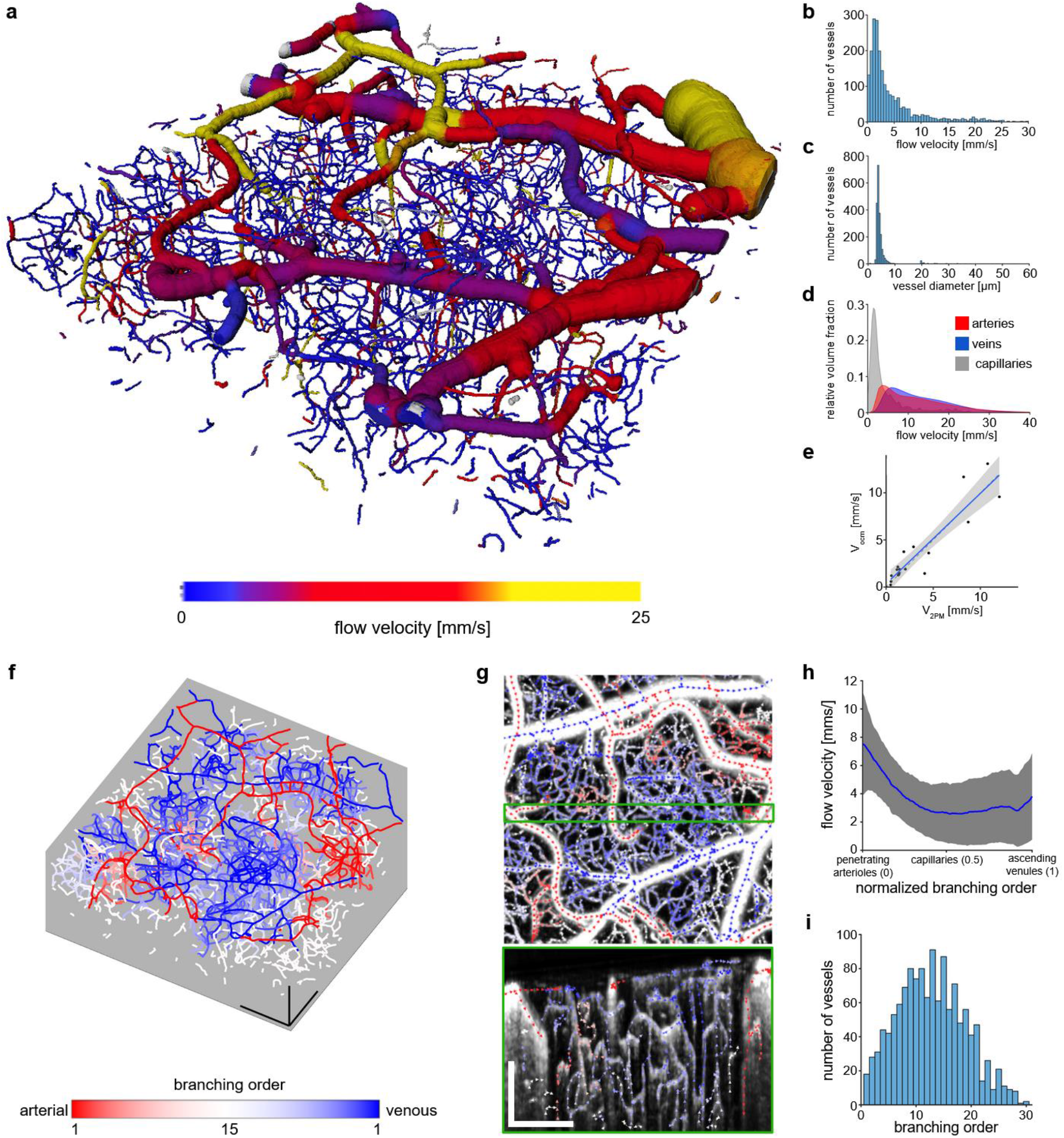
Vascular network analysis. a) Volumetric rendering of a fully segmented vascular network of size 1000 × 1000 × 360 μm^3^. The blood flow velocities in each segment are overlayed. b) Histogram of the flow velocities per vessel segment. c) Histogram of the average diameters per vessel segment. d) Flow velocities relative to the perfused volume for arteries, veins and capillaries. The relative volume fractions are computed only with respect to each vessel type. e) Velocity validation with respect to 2PM. The same animal was imaged sequentially with 2PM and OCM. 20 vessels were selected in the 2PM measurement and RBC flow velocity measured using line-scanning. The blue line indicates the linear fit (R^2^ = 0.84) and the shaded blue area the 95% confidence interval (CI: [0.63,0.9]) f) Graph of the branching order to the closest labeled vessel of the pial vasculature. Red indicates greater proximity to a pial artery, blue to a pial vein. Scalebars: 200 μm. g) Expanded maximum-intensity-projection (MIP) view of an angiogram over the first 50 μm of depth, overlayed with arrows indicating the flow direction. Arrow color indicates branching order as given in (f). A 40 μm thick side-view MIP is provided in the green box. Scalebars: 200 μm. h) Median velocity profile over all normalized shortest paths from the penetrating arterioles (0), through the capillary bed (0.5), to the ascending venules (1). For each vessel segment, the shortest path to any vessel segment labeled as an artery and any vessel segment labeled as a vein was extracted and traversed from the arterial to the venous side. Path length normalization is necessary to allow comparison and computation of the median value from all paths. The gray shaded area depicts the median absolute deviation (MAD). i) Histogram of the total branching order or equivalently path length of all paths used in h), before path length normalization.

In order to validate the proposed method *in vivo*, we sequentially performed imaging with 2PM and OCM. We chose 20 vessels and measured their respective RBC flow using 2PM. Afterwards, the same vessels were manually identified in the OCTA volume. Four vessels had to be excluded from comparison since strong tail artifacts from large pial vessels above hindered a successful segmentation. Higher blood flow velocities could not be compared due to the scanning speed limitations of the 2PM system. We find good agreement between the velocity measurements (Fig. 4e). Remaining differences are likely attributed to physiological changes over time, differing anesthesia depth or imperfect fits of the extracted Doppler spectra.

Accurate vessel classification into arteries and veins solely based on morphology is a challenging and time-consuming task that requires expert domain knowledge. It is further complicated when considering microscopes featuring capillary-level resolution where the FOV is usually limited with larger vessels not easily traceable back to known landmarks that could aid their classification. However, in the pial vasculature, the blood flow direction provides a clear indication of the vessel type. In arteries, the flow is directed towards smaller branches, distributing oxygen rich blood to the brain via diving arterioles. In contrast, the flow in veins is in the direction of larger branches, collecting de-oxygenated blood from ascending venules into larger veins. We further exploited the directional flow information in conjunction with vascular connectivity and morphology to facilitate efficient manual labeling of the pial vasculature. By projecting vessel-wise flow directions onto an angiogram maximum-intensity-projection (MIP) (Fig. S3), manual artery/vein classification for a 1000 × 1000 μm^2^ lateral FOV can be accomplished in less than 10 minutes.

From the labeled pial vasculature, the remaining vessel segments in the network are assigned via label propagation until reaching pre-capillaries, which are identified as having a median diameter below 12 μm. Furthermore, each vessel segment is assigned an arterial and a venous branching order by tracing the shortest path to the closest segment belonging to the respective pial vasculature trees (Fig. 4f). Fig. 4g displays a zoomed-in MIP view of an angiogram from Fig. 4f, in which the arrows indicate flow direction and their colors correspond to the branching order. Branches of a pial artery can be identified by observing the flow moving from the upper left corner toward the central part of the images. The branches of a vein are predominantly oriented horizontally. The green-framed side-view MIP accentuates the variations in the flow direction and vessel branching orders across the entire imaging depth. It also illustrates how the diving arterioles and ascending venules supply/collect blood to/from the deeper capillary bed.

Finally, we investigated how the flow velocities change throughout the network when moving from arteries to veins. For each vessel segment not classified as artery or vein, the shortest paths to the closest artery and vein are computed. Subsequently, the flow velocities along the combined path from the artery to the vein are gathered. In order to reject falsely allocated direct connections between arteries and veins, only paths that include at least three capillary segments are considered. All paths in the network are normalized to unit length to allow for a direct comparison. Median velocity of the capillary bed reaches its peak in proximity to diving arterioles, then dropping below 3 mm/s toward the center of the capillary bed (Fig. 4h). The velocities rise again towards ascending venules but remain well below the arterial velocities. The comparatively elevated median velocities in the capillary bed can be ascribed to the path length normalization, which may weight short connections between arteries and veins with higher velocities against connecting paths that traverse deeply through the capillary bed and display slow flow velocities. In order to account for the path length normalization, Fig. 4i shows a histogram of all path lengths before normalization. The capillary bed is characterized by an increasing number of vessels at higher branching orders until a branching order of approximately 12, forming an intricate network. The branching order distribution follows an approximately Gaussian shape until a branching order of 25.

## DISCUSSION

In this study, we introduce an advanced OCM approach that employs Bessel beam illumination for extended focus alongside a dedicated data-processing pipeline. The newly introduced framework is shown capable of analyzing morphological and functional parameters of the vascular network across multiple scales and particularized vessel types, enabling comprehensive examination of both network-wide effects and the intricate contributions of individual vessels, all the way from pial arteries and veins to the finest capillaries. The system’s extended focus offers an expansive field-of-view (1000 × 1000 × 360 μm^3^), allowing for snapshot acquisitions capturing the characteristics of large vascular networks comprehensively. To address the challenge of analyzing complex, information-dense datasets, we implemented automatic tools, including a deep learning-based 3D vessel segmentation network paired with highly sensitive DOCT for extraction of total flow velocity and direction. This approach not only highlights the importance of data-driven research in advancing our understanding of vascular networks but also demonstrates unprecedented sensitivity to flow dynamics, facilitating novel insights that cannot be carried out using a manual analysis.

Thus far, *in vivo* studies of the microvascular network’s morphology and function have often been performed with a combination of LSCI and 2PM^3,43^. Our method provides the high level of detail approaching that of 2PM while offering morphological information and total flow velocity measurements across all imaged vessels. Notably, a recent OCM study has looked into longitudinal vascular network alterations during the onset of Alzheimer’s disease^7^. However, conventional OCM approaches are plagued by hard compromises between the spatial resolution, depth of field and imaging speed, which limits their ability to extract and quantify vascular network characteristics across scales. A multi-modality approach has also been introduced, exploiting the superior resolution and SNR of 2PM combined with traditional DOCT to probe into flow velocities^44^. In contrast, OCTA volumes are directly segmented with our method, thus circumventing the higher system and registration complexity of the multi-modality system. Furthermore, our approach does not rely on angle estimation, which is inherently noisy when it comes to laterally branching vessels commonly found in the capillary bed. The same applies to the primarily laterally oriented pial vasculature for which even small angle estimation errors may lead to severely over-or under-estimated total flow velocities.

In summary, the introduced comprehensive, data-driven xf-irOCM pipeline represents a significant leap forward in *in vivo* quantification of morpho-functional parameters of cerebral vasculature across scales. Our approach offers new perspective on the microvascular connectome of the living brain, thus extending the frontiers of research into cerebrovascular function and neurovascular pathologies.

## METHODS

### xf-irOCM system

The imaging setup is devised according to the general concept of extended-focus OCM design^24,25^. A number of adjustments have been made in order to facilitate optimal velocity estimation. Briefly, at the system’s core is a Mach-Zehnder interferometer that allows splitting the illumination and detection modes. The xf-irOCM employs a broad-bandwidth SLED source (EBS300040-02, Exalos AG, Switzerland), centered at 1315 nm. The light source has a 3dB optical bandwidth of 120 nm and an output power of 30 mW. The light is split by a 90:10 beamsplitter and subsequently guided into the imaged sample and reference arms, respectively, optimizing the available power on the sample with respect to the previous iteration.

The light in the sample arm is similarly coupled into free space and then guided through a 178-degree axicon (Asphericon, Germany) to synthesize a Bessel beam. The beam is spatially filtered using a custom mask, rejecting stray light from the center of the annular beam. A pierced mirror decouples the illumination and detection light paths. Beam scanning is performed using two galvanometer scanners. A 10x IR-optimized objective (LMPLN10XIR, Olympus, Germany) focuses the light on the sample. The back-scattered light is collected via Gaussian mode detection as originally proposed by Leitgeb et al.^25^. Critically, the collected light’s path crosses the pierced mirror through the central hole and is then coupled into a single-mode fiber to be injected into a custom-built spectrometer for detection by a line scan camera (SU1024LDH2, Sensors Unlimited, USA).

### OCM data acquisition

In this work, two different acquisition protocols are used for OCTA and DOCT. After anesthesia and prior to imaging, a 100 μl bolus injection of 20 % Intralipid is administered. The scanned FOV is kept constant at 1 mm × 1 mm in both cases. For the OCTA acquisition, an A-scan rate of 46 kHz is used. Along the fast axis, 8 repeated B-scans with 512 pixels are recorded at each position. This is repeated for each of the 512 positions along the slow axis, resulting in an isotropic lateral pixel size of approximately 2 μm. The DOCT acquisition protocol uses an A-scan rate of 23 kHz and M-mode scanning, acquiring 250 A-scans at each lateral position, before moving to the next one. Here, the scanning grid consists of 256 pixels along both lateral dimensions, yielding a pixel size of 4 μm. The total acquisition time for one DOCT volume is approximately 12 minutes. In this work, three DOCT volumes are acquired for averaging.

### Image registration

The Doppler volume is registered to the angiogram volume from the same acquisition. This step is necessary due to the different scanning regimes. Instead of using the Doppler velocity volume directly, we use a volume in which each voxel corresponds to the maximum value of its respective Doppler spectrum (Doppler intensity). This resembles an angiogram the most after high-pass filtering. First, we fit a 3D affine transform, followed by a 3D deformable transform with 6 × 6 × 6 anchor points.

### High-sensitivity velocity estimation from Doppler OCT

We use the joint spectral-time domain OCT (jSTdOCT) framework to facilitate Doppler signal detection with high sensitivity^27^. Briefly, at each spatial location the 250 reconstructed A-scans are treated as a timeseries. First, a high-pass filter with a cut-off frequency of 270 Hz is employed along the time dimension. Afterwards, a Fourier transform, zero padded to 4096 values, is applied along the time dimension. This, following the jSTdOCT approach, yields interpolated Doppler spectra for each voxel. Subsequently, a modified Gaussian is independently fit to each Doppler spectrum as described previously by Bouwens et al.^28^. Due to the large number of necessary fits, we take advantage of the Gpufit library to provide fast fitting performance at large scale^45^. Additionally, the resulting fit parameters and goodness of fit (R^2^) are used to filter out badly fitted spectra and static tissue voxels. An additional step further rejects remaining static tissue voxels by adaptive thresholding of the Doppler spectrum intensity volume. Finally, axial and transverse velocity components are calculated for the remaining Doppler spectra, using mean and standard deviation, respectively^28^.

### Blood vessel segmentation and noise reduction

To accurately segment blood vessels at all scales in OCTA volumes, we employ the U-Net architecture^38^, which has been shown to achieve high-quality segmentation maps in various 2D and 3D biomedical imaging scenarios^39^. The detailed architecture of our 3D U-Net variant is depicted in Fig. S4.

To mitigate the need for large-scale manual annotations, we first pre-train our U-Net for 300 epochs on synthetic data. In this context, the synthetic images and their corresponding ground truth labels originate from physiologically plausible vascular trees that have been constructed based on a simplified angiogenesis model^40^. While vascular trees transformed to 3D volumetric data represent our ground truth labels, synthetic images rely on adjusting fore- and background intensity patterns to OCTA images at hand^36^. The resolution of the synthetic data has been adjusted so that small, simulated vessel segments match the size of capillaries in our OCTA images.

Subsequently, we finetune the U-Net for 100 epochs on manually annotated data. The annotation process was conducted and validated by two trained experts, who manually labeled a volume of 160 × 160 × 160 voxels (350 × 350 × 350 μm^3^). During the annotation process, special care was given to exclude tail artifacts and account for blood-flow-dependent variations of vessel signal intensities^46^. The volume selected for manual annotation contains an equal portion of large pial vessels and capillary bed and can, therefore, be seen as highly representative of the characteristics of OCTA volumes analyzed in this work. We find finetuning on manually annotated data to be a crucial step to adequately cope with OCTA-specific artifacts^33,47^.

We trained our 3D U-Net variant on a single NVIDIA V100 GPU (16 GB) using the AdamW optimizer with a learning rate of 1e^-5^, optimizing the average Dice loss function^48^, which is particularly suited for segmentation tasks subject to class imbalance. During training, we feed mini-batches of size 32, composed of image crops at random locations of the shape of 64 × 64 × 64 voxels, to the segmentation model. With a batch size of 32, each epoch consists of 312 iterations. To increase the diversity of our training data and hence tackle overfitting, we make use of several data augmentation techniques. Specifically, we apply a variety of intensity-based (random Gaussian noise, random Gaussian smoothing, random transformations to the image’s intensity histogram) and spatial (randomly flipping, random rotation, random zoom) augmentations. Since our U-Net has been trained on crops, we employ a sliding window inference scheme with an overlap of 90 % between crops.

### Skeletonization and graph construction

Skeletonization and graph construction is performed using Voreen^41^. It performs well on vessel datasets, rendering subsequent skeleton cleaning unnecessary. Voreen only requires the non-dimensional input *bulge size* which we empirically optimize to 3. The graph representation assigns nodes to all skeleton bifurcation- and endpoints while edges represent the remaining skeleton. Each edge is also assigned further information such as length, diameter, tortuosity among others. Furthermore, for each edge, one connected node is arbitrarily assigned to be the start and the other to be the endpoint.

### Vessel diameter fitting

We ensure accurate vessel diameters at capillary scale through an additional cross-section fitting step. This is performed along all vascular network skeleton voxels. At each skeleton voxel, a 1D Gaussian function is fit to a line profile obtained from the OCTA volume. The line profile is orthogonal to the vessel direction to obtain more accurate vessel diameters. Only fitted Gaussians with an R^2^ > 0.8 are kept. Then, a moving average is used to obtain the diameter profile along the vessel segment and an average diameter for the entire vessel is calculated.

### Velocity estimations in individual segments

Vessel segment-wise velocity data is obtained by combining the registered total flow velocity Doppler volumes and locally averaging spheres with diameters extracted during the previous fitting step. For each skeleton voxel, we gather all total as well as axial flow velocity values. Then, the skeleton position is assigned total and axial velocity values computed as the mean of the highest 30 % of respective velocity values within the local sphere. Next, a moving average operation is performed along each vessel segment’s skeleton voxels to ensure smooth velocity variation along the vessel segment. Finally, we compute the total flow velocity for the vessel segment as the median value of all skeleton positions.

### Estimation of the missing velocities

In case of missing velocities, we estimate the segment’s flow velocity from its connected vessel segments (Fig. S5). At each of its two nodes, we enforce conservation of mass, requiring equal inflow and outflow, that is equivalent to a zero net flow at each node. Here, we take advantage of known flow directions and assume that the local blood density is constant. Using the flow information from the other connected vessels and the known vessel diameter, we estimate an average flow velocity as the average of the computed value from its two nodes. If one node is an end node or has multiple connected vessels with missing velocity information, only the other fully described node is considered. This algorithm is iteratively repeated until all segments have been assigned a velocity or no more segments can be fixed.

### Phantom experiments

We performed multiple phantom experiments to validate our method for total blood flow velocity estimation. All phantom experiments are conducted using 180 μm inner diameter plastic tubing. We used 1.5 % intralipid solution as a blood substitute. A syringe pump was employed to accurately control flow velocities. To evaluate the accuracy of the flow velocities, the tubing is placed at an 8-degree angle and the velocity is slowly increased in steps with ample time in between to ensure steady-state syringe pump performance (Fig. S1). In a second phantom experiment, the syringe pump is set to a constant rate, resulting in a flow velocity of 10 mm/s. Then, the phantom is axially shifted through the focus to validate out-of-focus performance (Fig. S2).

### Animal handing

Experiments were performed on female C57Bl6/J mice (no.028, Charles River Laboratories, USA), 6 to 12 weeks of age, weighing 25 - 30 g. The mice were housed under standard conditions, including free access to water and food as well as an inverted 12-hour light/dark cycle. For headpost and cranial window implantation, animals were injected intraperitoneally with a triple mixture of fentanyl (0.05 mg/kg body weight; Sintenyl, Sintetica, Switzerland), midazolam (5 mg/kg body weight; Dormicum, Roche, Switzerland) and medetomidine (0.5 mg/kg body weight; Domitor, Orion Pharmaceuticals, Switzerland). The face mask delivers 100 % oxygen at a rate of 300 mL/min. After cranial window implantation, mice were allowed to recover for two weeks prior to imaging. For OCT and 2PM imaging, mice received the same triple anesthesia mixture intraperitoneally. During all procedures, the animals’ core temperature was maintained at 37 °C using a thermostatic blanket heating system (Harvard Apparatus, USA). All animal experiments were performed in accordance with the Swiss Federal Act on Animal Protection and approved by the Cantonal Veterinary Office Zurich.

### Two-photon microscopy reference experiments

Two-photon microscopy (2PM) was performed to obtain reference velocity measurements in-vivo. After anesthesia, Cascade blue Dextran (5 % w/v, 10,000 kDamw, 100 μl, D-1976, Life Technologies, USA) was injected intravenously 10 minutes before imaging and excited at 820 nm to visualize the vasculature. Next, we acquired a 300 μm depth-stack of 400 × 400 μm^2^ images of the cortical vasculature. From the 3D image stack, we chose 20 vessels and measured their respective RBC velocities using repeated line scanning and the Radon algorithm^49^. Subsequently, we acquired an OCM dataset at the same position using the method described in this work. Finally, we manually matched the 20 vessels between the two datasets. Four of the vessels could not be matched because they were obscured by the tail artifacts of larger vessels in the OCM dataset.

### Software and statistical analysis

The processing pipeline was developed using Matlab (R2021b, MathWork, USA). Additionally, we used some functions from the open-source code provided by Stefan et al.^42^. Registration was done using SimpleITK (v2.3, simpleITK.org). The deep learning pipeline builds upon PyTorch (pytorch.org) and MONAI (monai.io). The 2PM was controlled by a customized version of ScanImage (r3.8.1, Janelia Research Campus). Analysis of the *ex vivo* vessels graphs from Blinder et al.^1^ was performed using Python (Python 3.10). Volumetric renders as shown in Fig. 1d and Fig. 4a were rendered using Amira (2020.3, Thermo Fisher Scientific Inc., USA). Plots were created using either Matlab (R2021b, MathWork, USA) or R (v4.3, r-project.org) and the ggplot2 package (v3.4, ggplot2.tidyverse.org).

## Acknowledgments

D.R. acknowledges funding from the Swiss National Science Foundation (310030_192757) and the US National Institutes of Health (R01-NS126102). B.M. is supported through a Helmut-Horten-Professorship for Biomedical Informatics by the Helmut Horten Foundation. B.Wi. is supported through the Helmut Horten Foundation. S.W. acknowledges funding from the Swiss National Science Foundation (310030_200703) and the UZH CRPP stroke.

The authors appreciate the help of Paul-James Marchand with setting up the OCM system as well as fruitful discussions and thank Sabina Stefan for discussions of her segmentation pipeline and sharing of her training data for comparison.

## Author Contributions Statement

L.G. and D.R. conceived the study. L.G. developed the experimental system. L.G., J.D. and C.G. carried out the animal experiments. L.G. and B.Wi. developed the processing pipeline and conducted the data analysis and visualization. J.D., C.G., M.E.A., R.L. and B.M. contributed to the interpretation of the results. B.We. and S.W. provided the animal model and guided the animal experiments. D.R, R.L. and B.M. were involved in the study design and planning and supervised the work. All authors contributed to writing and revising the manuscript.

## Data Availability

The data that support the findings of this study are available from the corresponding author upon reasonable request.

