## supplemental figures for "Bessel Beam Optical Coherence Microscopy Enables Multiscale Assessment of Cerebrovascular Network Morphology and Function"

**Supplementary Figure 1:**
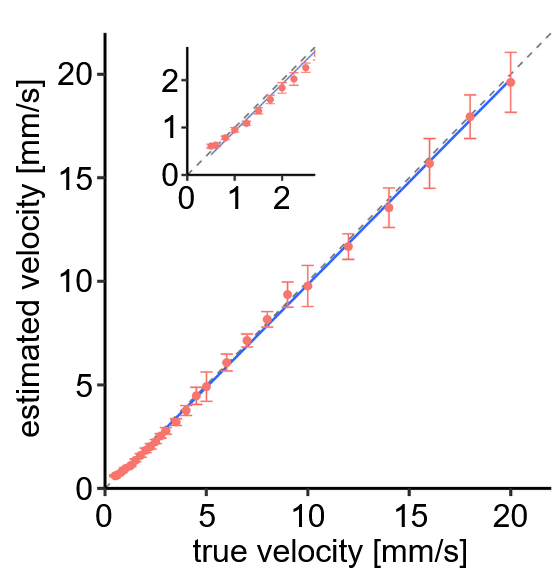

Figure 1: Phantom study of velocity measurement in 180 μm tubing and 1.5% intralipid solution. Good agreement between the ground truth and estimated flow velocities is found. Flow velocities above ≈20 mm/s result in Doppler aliasing due to the 8° angle of the tubing but do not constitute an upper limit for velocity estimation in-vivo. The pial vasculature, exhibiting the highest flow velocities, typically has angles closer to 0°, preventing aliasing.

**Supplementary Figure 2:**

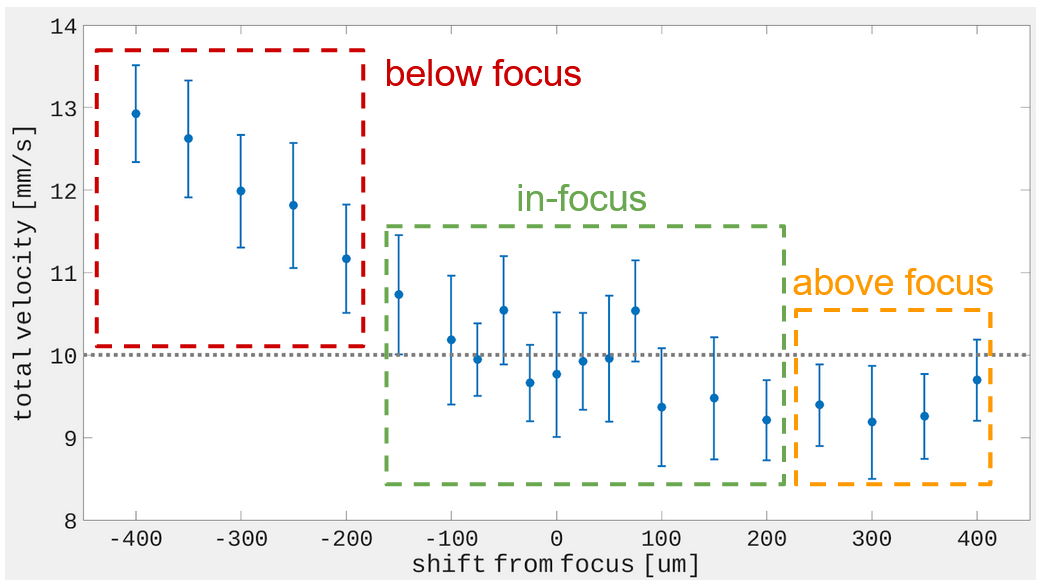

Figure 2: Phantom study of out-of-focus velocity estimation in 180 μm tubing and 1.5% intralipid solution. Flow well above the focus is slightly underestimated (<10%). Only flow far below the in-focus volume showed flow overestimation in excess of 10%. For in-vivo studies, this would not be considered because the microvasculature is not segmented in the regions far below the in-focus volume.

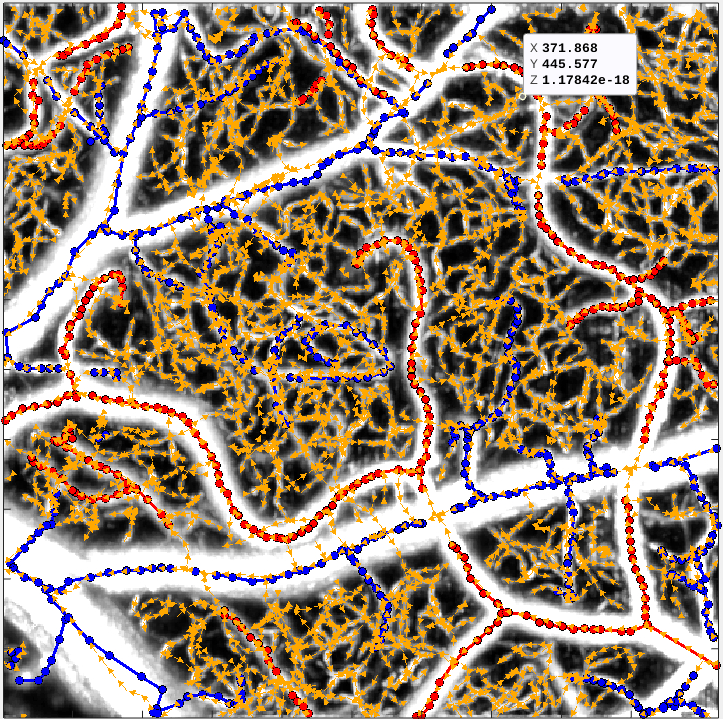
**Supplementary Figure 3:**

**veins**

**arteries**

Figure 3: Angiogram MIP with overlayed flow directions (yellow) and manual artery (red) and vein (blue) labels.

**Supplementary Figure 4:**
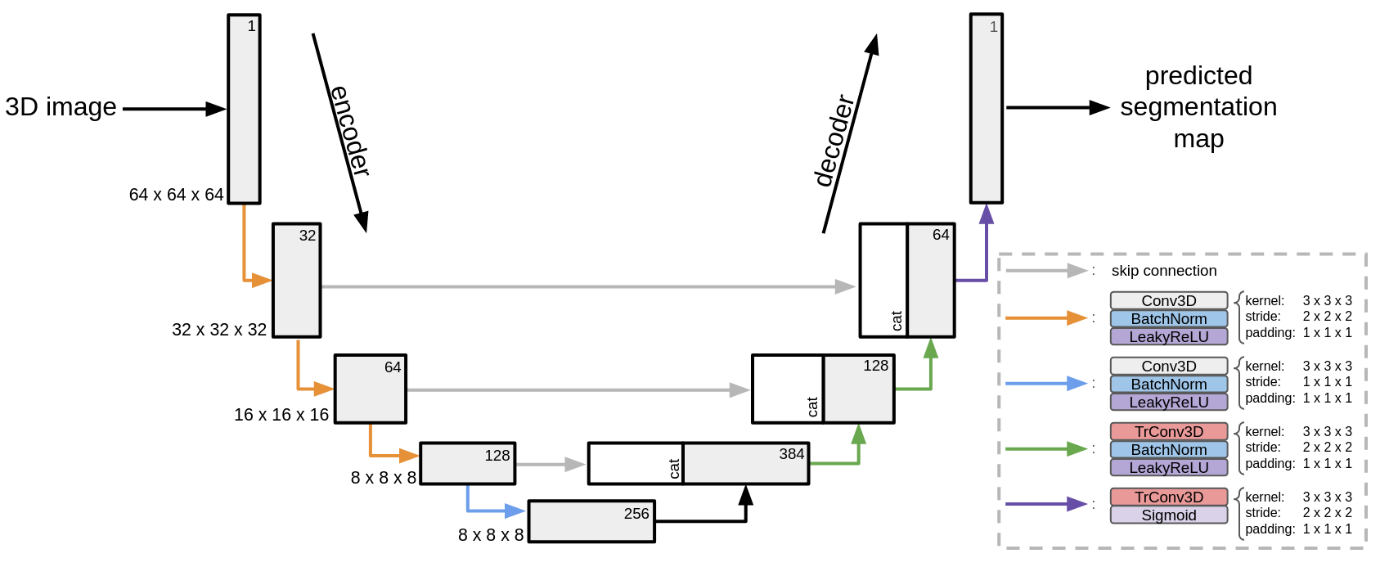

Figure 4: Detailed architecture of our employed 3D U-Net. We would like to draw the reader’s attention to color coding.

**Supplementary Figure**
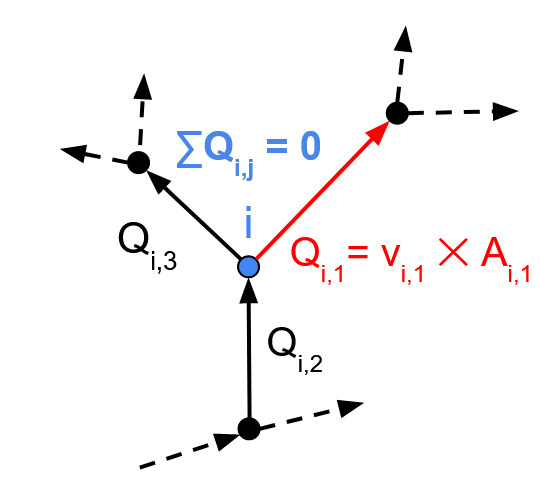
**5:**

Figure 5: Estimating missing velocities through conservation of mass at branch points.
